# Isolation of protein N-terminal peptides by charge-mediated position-selective enrichment using strong cation exchange chromatography

**DOI:** 10.1101/2020.06.15.153684

**Authors:** Chih-Hsiang Chang, Hsin-Yi Chang, Juri Rappsilber, Yasushi Ishihama

## Abstract

We developed a simple and rapid method to enrich protein N-terminal peptides, in which the protease TrypN is first employed to generate protein N-terminal peptides without Lys or Arg and internal peptides with two positive charges at their N-termini, and then the N-terminal peptides are separated from the internal peptides by means of CHArge-Mediated Position-selective (CHAMP) enrichment using strong cation exchange (SCX) chromatography. This CHAMP-SCX approach was applied to 20 μg of human HEK293T cell lysate proteins to profile the N-terminal proteome. On average, 1,550 acetylated and 200 unmodified protein N-terminal peptides were successfully identified in a single LC/MS/MS run with less than 3% contamination with internal peptides, even when we accepted only canonical protein N-termini registered in the Swiss-Prot database. Since this method involves only two steps, protein digestion and chromatographic separation, without the need for tedious chemical reactions, it should be useful for comprehensive profiling of protein N-termini, including proteoforms with neo-N-termini.

## INTRODUCTION

Characterizing protein N-termini is essential to understand how the entire proteome is generated through biological processes such as translational initiation (1-3), post-translational modifications (4, 5) and proteolytic cleavages (6, 7). In order to perform N-terminomics using mass spectrometry (MS), peptides derived from protein N-termini must be selectively enriched, and many methods have been developed for this purpose (8, 9). Some of them use “positive selection” approaches in which chemically labeled protein N-terminal peptides are enriched by affinity purification (6, 10). However, these approaches are difficult to apply to protein N-termini with modifications. In contrast, “negative selection” approaches to isolate protein N-terminal peptides by depleting internal peptides have been used to comprehensively identify protein N-terminal peptides, including N-terminal modifications such as methylation, acetylation, and lipidation (11, 12). Gevaert *et al.* pioneered combined fractional diagonal chromatography (COFRADIC) (13), and this was followed by other negative selection approaches such as terminal amine isotopic labeling of substrates (TAILS) (14), the variant of COFRADIC called charge-based fractional diagonal chromatography (ChaFRADIC) (15), and hydrophobic tagging-assisted N-termini enrichment (HYTANE) (16). All of them require blocking of the primary amines at the protein level and depletion of digested internal peptides by means of chemical tagging-based separation. Thus, relatively large amounts of samples (∼5 to 10 mg) are generally required to increase the identification number of protein N-terminal peptides. This limits the usefulness of these approaches in the case of hard-to-obtain biological samples (17, 18). Furthermore, limitations in the efficiency and specificity of the chemical derivatizations compromise the confidence of peptide identification. Therefore, a simple and sensitive approach to enrich protein N-terminal peptides is still needed for MS-based proteomics.

Strong cation exchange (SCX) chromatography, employing Coulombic interactions to separate peptides based on their positive charges, has been widely applied for deep proteomic profiling (19, 20). In SCX separation of tryptic peptides, abundant acetylated protein N-terminal peptides are eluted first. Peptides with 1+ charge such as monophosphorylated peptides, N-pyroglutamated peptides and protein C-terminal peptides are then simultaneously eluted, followed by peptides with +2 or more net charge, such as unmodified protein N-terminal peptides, internal peptides and peptides containing missed cleavages (21, 22). Thus, it is impossible to isolate protein N-terminal peptides from tryptic peptides by SCX chromatography. To overcome this issue, we focused on TrypN (23), also known as LysargiNase, a metalloprotease that cleaves peptide chains mainly at the N-terminal side of Lys/Arg even in the case of Pro-Lys and Pro-Arg bonds, generating peptides with N-terminal Lys/Arg and yielding protein N-terminal peptides that do not contain Lys/Arg. Unlike other LysargiNases such as ulilysin (24, 25) and mirolysin (26), which preferentially cleave the N-terminal side of either Lys or Arg, TrypN cleaves the N-terminal side of Lys and Arg equally at pH 6∼8. Moreover, the peptide identification performance for N-terminal Lys/Arg peptides is comparable to that for tryptic peptides (27).

In this study, we developed a new method to enrich protein N-terminal peptides without the need for chemical derivatization or complex procedures, taking advantage of the combination of proteinase TrypN-mediated protein cleavage and SCX separation of N-terminal peptides based on the charge difference at the N-termini of protein N-terminal peptides and internal peptides. We call this method CHArge-Mediated Position-selective (CHAMP) enrichment using strong cation exchange (SCX) chromatography (CHAMP-SCX). We show that this rapid and simple approach to enrich protein N-terminal peptides enables comprehensive, high-throughput analysis of the human N-terminal proteome.

## MATERIALS AND METHODS

### Materials

Ammonium bicarbonate, 2-amino-2-(hydroxymethyl)-1,3-propanediol hydrochloride (Tris-HCl), sodium deoxycholate (SDC), sodium N-lauroylsarcosinate (SLS), ammonium bicarbonate, tris(2-carboxyethyl)phosphine (TCEP), 2-chloroacetamide (CAA), calcium chloride, ethyl acetate, acetonitrile (ACN), acetic acid, trifluoroacetic acid (TFA) and other chemicals were purchased from Fujifilm Wako (Osaka, Japan). Modified trypsin was from Promega (Madison, MA). TrypN was from Protifi (Huntington, NY). Styrene divinylbenzene (SDB-XC) Empore™ disk and strong cation exchange (Cation-SR) Empore™ disks were purchased from GL Sciences (Tokyo, Japan). Water was purified by a Millipore Milli-Q system (Bedfold, MA, USA).

### *Escherichia coli* cell culture and protein extraction

*Escherichia coli* K-12 BW25113 was grown to mid-log phase in LB broth with vigorous shaking at 37°C. Cells were collected by centrifugation and resuspended in buffer containing protease inhibitors (Sigma), 12 mM SDC, 12 mM SLS, 10 mM TCEP, 40 mM CAA in 100 mM Tris buffer (pH 8.5). The lysate was vortexed and sonicated on ice for 20 min. The concentration of protein crude extract was determined by means of bicinchoninic acid (BCA) protein assay (Thermofisher Scientific, Rockford, IL, USA).

### HEK293T cell culture and protein extraction

HEK293T (human embryonic kidney) cells, cultured to 80% confluence in 10-cm diameter dishes, were harvested in lysis buffer containing protease inhibitors (Sigma), 12 mM SDC, 12 mM SLS, 10 mM TCEP, 40 mM CAA in 100 mM Tris buffer (pH 8.5). The lysate was vortexed and sonicated on ice for 20 min. The final protein concentration of the sample was determined using the BCA protein assay.

### Protein Digestion

For TrypN digestion, the quantified protein solution was diluted 10-fold with 10 mM CaCl2 and digested with TrypN (1: 50 w/w) overnight at 37 °C. Note that TrypN can be replaced with LysargiNase (Merck Millipore, Darmstadt, Germany). In the case of tryptic digestion, the protein solution was digested with Lys-C (1:50 w/w) for 3 h at 37 °C, followed by 5-fold dilution with 50 mM ammonium bicarbonate and trypsin digestion (1:50 w/w) overnight at 37 °C. After enzymatic digestion, an equal volume of ethyl acetate was added to each sample solution, and the mixture was acidified with 0.5% trifluoroacetic acid (final concentration) according to the PTS protocol reported previously (28). The resulting mixture was shaken for 1 min and centrifuged at 15,700 g for 2 min to separate the ethyl acetate layer. The aqueous layer was collected and desalted by using StageTips with SDB-XC disk membranes (29). The proteolytic peptides were quantified by LC-UV at 214 nm with standard BSA peptides and kept in 80% ACN and 0.5% TFA at −20 °C until use.

### Peptide fractionation by SCX HPLC

SCX chromatography was performed using a Prominence HPLC system (Shimadzu, Kyoto, Japan) with a BioIEX SCX column (250 mm × 4.6 mm inner diameter, 5 μm non-porous beads made of poly(styrene-divinylbenzene) modified with sulfonate groups (Agilent, Santa Clara, CA, USA).

For examination of the SCX separation characteristics, 75 μg each of trypsin- and TrypN-digested HEK293T peptides were mixed and directly loaded onto the SCX column at 0.8 mL/min. A mixture of 5 mM potassium phosphate (pH 3.0) and ACN (7:3) was used as SCX buffer A, and a mixture of 500 mM potassium phosphate (pH 3.0) and ACN (7:3) was used as SCX buffer B. Gradient elution was performed as follows: 0% B for 5 min, 0-10% in 20 min, 10-50% in 10 min, 50-100% in 5 min and 100% B for 4 min. Fractions were manually collected at one min intervals for 45 min. After evaporation of the solvent in a SpeedVac SPD121P (Thermofisher Scientific), fractionated peptides were resuspended in 50 μL of 0.1% TFA and desalted by using StageTips with SDB-XC disk membranes. One-fourth of each fraction was analyzed by nanoLC/MS/MS using a TripleTOF 5600 (SCIEX, Foster City, CA, USA) as described below.

For gradient SCX fractionation of TrypN-digested HEK293T peptides, 80 μg of digested peptides were analyzed using the SCX HPLC system described above. A mixture of 7.5 mM potassium phosphate (pH 2.6) and ACN (7:3) was used as SCX buffer A, and 350 mM KCl was added to buffer A for SCX buffer B. Gradient elution was performed as follows: 0.5% B for 15 min, 0.5-1% B in 10 min, 1-4% B in 10 min, 4-10% B in 3 min, 10-100% B in 3 min, and 100% B for 5 min. Fractions were manually collected at one min intervals for 50 min. After evaporation of the solvent in a SpeedVac, fractionated peptides were resuspended in 50 μL of 0.1% TFA and desalted by using StageTips with SDB-XC disk membranes. One-fourth of each fraction for Fr.1-43 and one-tenth of each fraction for Fr.44-50 were analyzed by nanoLC/MS/MS using an Orbitrap Fusion Lumos mass spectrometer (Thermofisher Scientific) as described below.

### Enrichment of protein N-terminal peptides by CHAMP-SCX

Enrichment of protein N-terminal peptides from 30 μg of TrypN-digested *E. coli* peptides was performed using the SCX HPLC system under the following isocratic conditions: SCX buffer A was a mixture of 7.5 mM potassium phosphate solution (pH 2.2) containing 10, 12.5 or 15 mM KCl and ACN (7:3), and buffer B was a mixture of 7.5 mM potassium phosphate solution (pH 2.2) containing 500 mM KCl and ACN (7:3). Isocratic elution was performed with 100% A for 30 min and then the system was washed with 100% B. The collected eluates were dried in a SpeedVac and the residue was resuspended in 50 μL of 0.1% TFA and desalted on StageTips with SDB-XC disk membranes. Two-thirds of the enriched peptides were analyzed by nanoLC/MS/MS using the Orbitrap Fusion Lumos.

To isolate protein N-terminal peptides from TrypN-digested HEK293T peptides, the digested peptides (80 μg) were analyzed by the SCX HPLC system under isocratic conditions, eluting with a mixture of 7.5 mM potassium phosphate solution (pH 2.2) containing 10 mM KCl and ACN (7:3) for 30 min to collect the desired fraction. After evaporation of the solvent in a SpeedVac, enriched peptides were resuspended in 50 μL of 0.1% TFA and desalted using StageTips with SDB-XC disk membranes. We analyzed one-fourth of the enriched peptides by nanoLC/MS/MS in triplicate using the Orbitrap Fusion Lumos.

### NanoLC/MS/MS analysis

NanoLC/MS/MS analyses were performed on a TripleTOF 5600 mass spectrometer or an Orbitrap Fusion LUMOS mass spectrometer, connected to a Thermo Ultimate 3000 pump and an HTC-PAL autosampler (CTC Analytics, Zwingen, Switzerland). Peptides were separated on self-pulled needle columns (150 mm length × 100 μm ID, 6 μm opening) packed with Reprosil-C18 3 μm reversed-phase material (Dr. Maisch, Ammerbuch, Germany). The injection volume was 5 μL and the flow rate was 500 nL/min. The mobile phases were (A) 0.5% acetic acid and (B) 0.5% acetic acid and 80% ACN. For TripleTOF 5600 analysis, gradient elution was performed as follows: 12−40% B in 20 min, 45-100% B in 1 min, 100% B for 5 min. For Orbitrap analysis, gradient elution of fractionated samples was performed as follows: 12−40% B in 15 min, 40-100% B in 1 min, 100% B for 5 min. For protein N-terminal peptide-enriched samples, gradient elution was performed as follows: 10−40% B in 100 min, 40-100% B in 10 min, 100% B for 10 min. Spray voltages of 2300 V in the TripleTOF 5600 system and 2400 V in the Orbitrap system were applied. The mass scan range of the TripleTOF 5600 system was *m/z* 300−1500, and the top ten precursor ions were selected in each MS scan for subsequent MS/MS scans. The mass scan range for the Orbitrap system was *m/z* 300−1500, with an automatic gain control value of 1.00e + 06, a maximum injection time of 50 ms and detection at a mass resolution of 60,000 at *m/z* 200 in the orbitrap analyzer. The top ten precursor ions were selected in each MS scan for subsequent MS/MS scans with an automatic gain control value of 5.00e + 04 and a maximum injection time of 300 ms. Dynamic exclusion was set for 25 s with a 10 ppm gate. The normalized HCD was set to be 30, with detection at a mass resolution of 15,000 at *m/z* 200 in the Orbitrap analyzer. A lock mass (445.1200025) function was used to obtain constant mass accuracy during the gradient.

### Proteomics Data Processing

Two peak lists in “.mgf” and “.apl” formats were generated from the MS/MS spectra by MaxQuant (30). The peptides and proteins were identified by Mascot v2.6.1 (Matrix Science, London, U.K.) against the Swiss-Prot database (version 2017_4, 20,199 sequences) or the *E. coli* K-12 MG1665 protein sequence database (31) with a precursor mass tolerance of 20 ppm (TripleTOF 5600) or 10 ppm (Orbitrap), a fragment ion mass tolerance of 0.1 Da (TripleTOF 5600) or 20 ppm (Orbitrap), TrypN/trypsin specificity allowing for up to 4 missed cleavages for TrypN/trypsin mixed proteolytic peptides and strict TrypN specificity allowing for up to 2 missed cleavages for TrypN-digested peptides. Carbamidomethylation of cysteine was set as a fixed modification, and methionine oxidation and protein N-terminal acetylation were allowed as variable modifications. False discovery rates at a peptide level of less than 1% were applied for peptide identification based on a target-decoy approach.

## RESULTS AND DISCUSSION

### Charge-mediated position-selective enrichment in SCX chromatography

Proteolysis with TrypN yields peptides with at least a +2 charge with Lys or Arg and an α-amino group at the N-terminus. On the other hand, peptides derived from protein N-termini have neither Lys nor Arg and are highly acetylated at the N-terminus, so that most of them have a 0 or +1 charge, and only His-containing peptides with an unmodified N-terminus have a +2 charge. In this study, we focused on the fact that SCX chromatography at low pH can separate peptides based on the number of positive charges, and we attempted to separate protein N-terminal peptides from internal peptides among TrypN-digested peptides.

We first examined the number of missed cleavages in TrypN digestion. When digestion was performed in 0.1% RapiGest according to the manufacturer’s protocol, the missed cleavage rate (the content of peptides with two or more missed cleavage sites) was 14%, almost equal to the value in the condition without addition of RapiGest (16%). On the other hand, when 1% SDC was added instead of RapiGest, the missed cleavage rate was dramatically reduced to 5.8%. Thus, TrypN digestion was performed according to the PTS protocol (28) in this study.

Next, keeping in mind the need to separate protein N-terminal-derived peptides with His residues and unmodified N-termini from TrypN-digested internal peptides, we investigated whether the peptides could be separated based on the position of the positive charge, in addition to the number of positive charges, by SCX chromatography. Studies with proteases that cleave either Lys/Arg, such as Lys-C and Lys-N or trypsin and TrypN, have indicated that the position of the positive charge affects the outcome in shotgun proteomics (25, 32). For example, it has been reported that peptides with N-terminal Lys or Lys/Arg are more strongly retained than peptides with a C-terminal Lys or Lys/Arg in reversed-phase LC (27). To determine how the Lys/Arg position of peptides affects their retention behavior in SCX chromatography at low pH, we examined a mixture of TrypN- and trypsin-digested peptides using the SCX HPLC system, followed by nanoLC/MS/MS. The 19,853 unique tryptic peptides generally showed weaker retention than the 11,334 unique TrypN peptides with the same charge states (Supplementary Fig. S1). To characterize the SCX elution profiles in more detail, we compared the retention time in SCX HPLC for approximately 4,000 peptide pairs having sequences that differ only in the presence or absence of terminal Lys/Arg (Figure 1A). As expected, a strong retention shift was observed for TrypN-digested peptides. This would be because the TrypN peptides are strongly positively charged at the N-terminus, due to the α-amino group of the N-terminal Lys/Arg and the side chain ε-amino or guanidino group, whereas the positive charge of the C-terminal Lys/Arg of trypsin peptides was partially neutralized by the α-carboxy group (Fig. 1B). Gauci *et al*. reported that Lys-N-digested phosphopeptides with two basic moieties in close proximity tend to be more strongly retained on an SCX column than tryptic phosphopeptides (22). Gussakovsky *et al.* reported a retention model for predicting the retention times in SCX chromatography of tryptic peptides, in which the position-dependent coefficient of basic amino acids is higher near the N-terminus (33). We also found that the TrypN-digested peptides eluted in a narrower SCX fraction range than the tryptic peptides (Fig. S1). This may be due to the fact that the N-terminal positive charge of the tryptic peptides differs depending upon the amino acid involved, whereas the N-terminal amino acid of the TrypN-digested peptide is always Lys or Arg. To our knowledge, the present work is the first to show that the position of basic residues in the same sequence reliably affects the electrostatic interaction with the SCX stationary phase using thousands of identical sequence pairs, and we named this approach charge-mediated position-selective SCX (CHAMP-SCX) enrichment.

**Figure 1.**
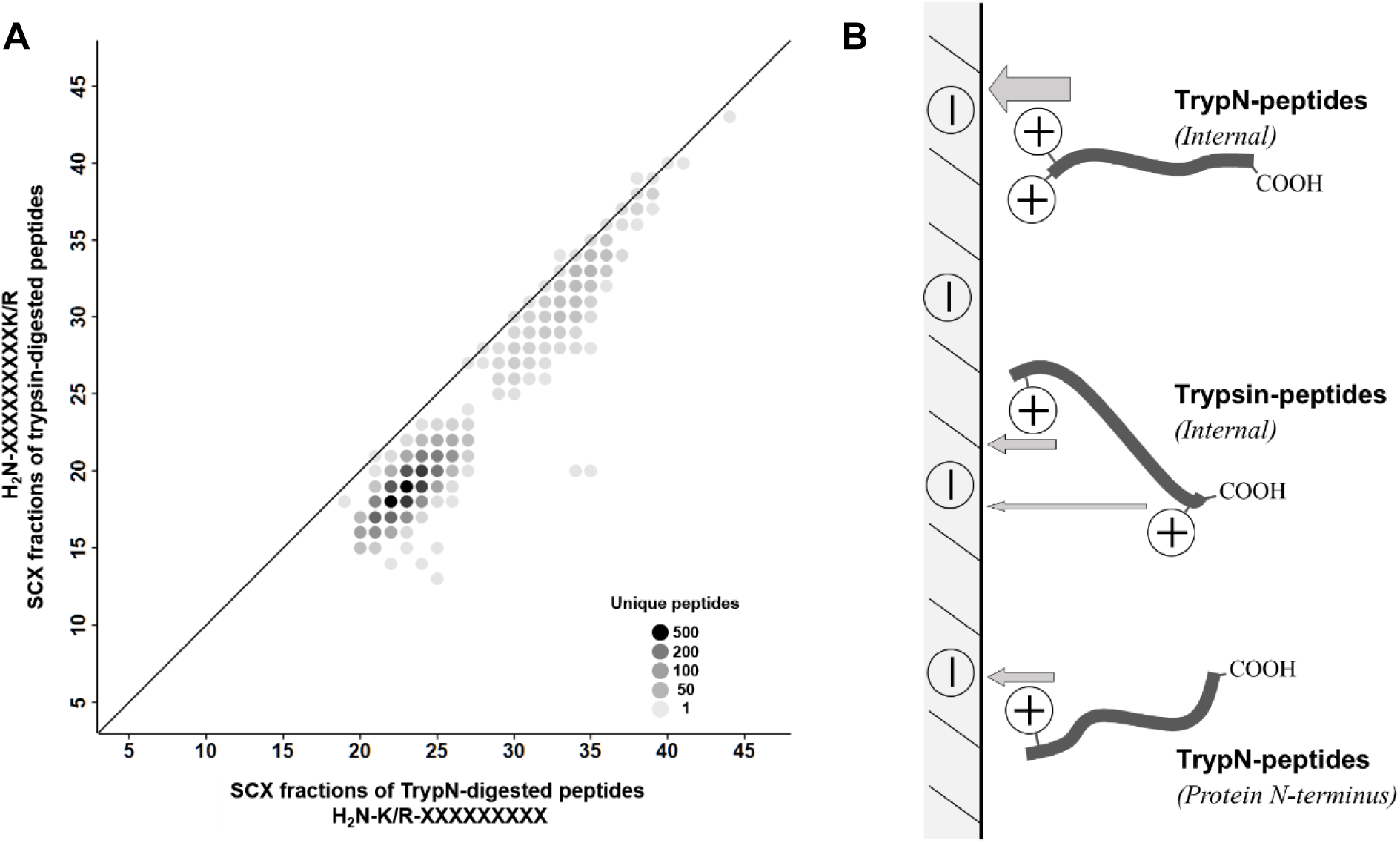
SCX separation of TrypN-digested and trypsin-digested peptides. (A) Comparison of SCX elution profiles of TrypN- and trypsin-digested peptides. Peptides with the same sequence except for the termini were selected (K/R-XXXXXX and XXXXXX-K/R for TrypN- and trypsin-digested peptides, respectively). The shade of the circle color indicates the number of peptides. (B) Proposed retention model of TrypN-digested internal peptides, trypsin-digested internal peptides and TrypN-digested protein N-terminal peptides on SCX stationary phase.

### CHAMP-SCX separation of TrypN-digested HEK293T peptides

The HPLC system used in this study was equipped with a nonporous hydrophilic SCX column having a separation efficiency equivalent to that of a typical reversed-phase column (the peak width at half height was 12.4 ± 4.2 seconds and the peak capacity was 122; see Supplementary Fig S2). As already shown in Supplementary Figure S1, this SCX HPLC system was able to separate TrypN-digested peptides with +1 and +2 charges from each other. Comprehensive SCX fractionation of TrypN-digested peptides derived from HEK293T cells was performed with a KCl salt gradient elution at pH 2.6, and peptide identification for each fraction was performed by nanoLC/MS/MS. As shown in Figure 2A, nearly all of the protein N-terminal-derived peptides were clearly separated from the internal peptides, regardless of whether their N-termini were acetylated or not. The fractions from 2 to 11 min contained mainly 0 and +1 peptides, including 2,207acetylated protein N-terminal peptides, 345 His-containing acetylated N-terminal peptides, and 262 unmodified N-terminal peptides. The 12-18 min fractions contained +2 peptides, *i.e.*, unmodified protein N-terminal peptides containing one His, Lys or Arg and acetylated protein N-terminal peptides containing two basic amino acids. The next fractions from 19 to 30 min also contained +2 peptides, but most of them were internal peptides based on the CHAMP effect, i.e., retention was stronger due to the high density of positive charge at the N-terminus of the peptides (Fig. 1B). Thus, the protein N-terminal peptides can be easily isolated. Peptides with a charge greater than 2+ were sequentially eluted in the fractions after 31 min. These included protein N-terminal peptides containing missed cleavage sites, but their number was small due to the high efficiency of TrypN digestion by the PTS method. Up to 90% of non-redundant protein N-terminal peptides could be recovered in fractions up to 18 min by CHAMP-SCX separation (Supplementary Figure S3), demonstrating that the combination of TrypN digestion and CHAMP-SCX enables simple and rapid protein N-terminal peptide enrichment. In addition, unlike trypsin, which is unable to cleave Lys-Pro and Arg-Pro bonds, TrypN can cleave Pro-Lys and Pro-Arg bonds, generating protein N-terminal peptides with Pro at the C-termini, and thus improving the coverage in N-terminomics.

**Figure 2.**
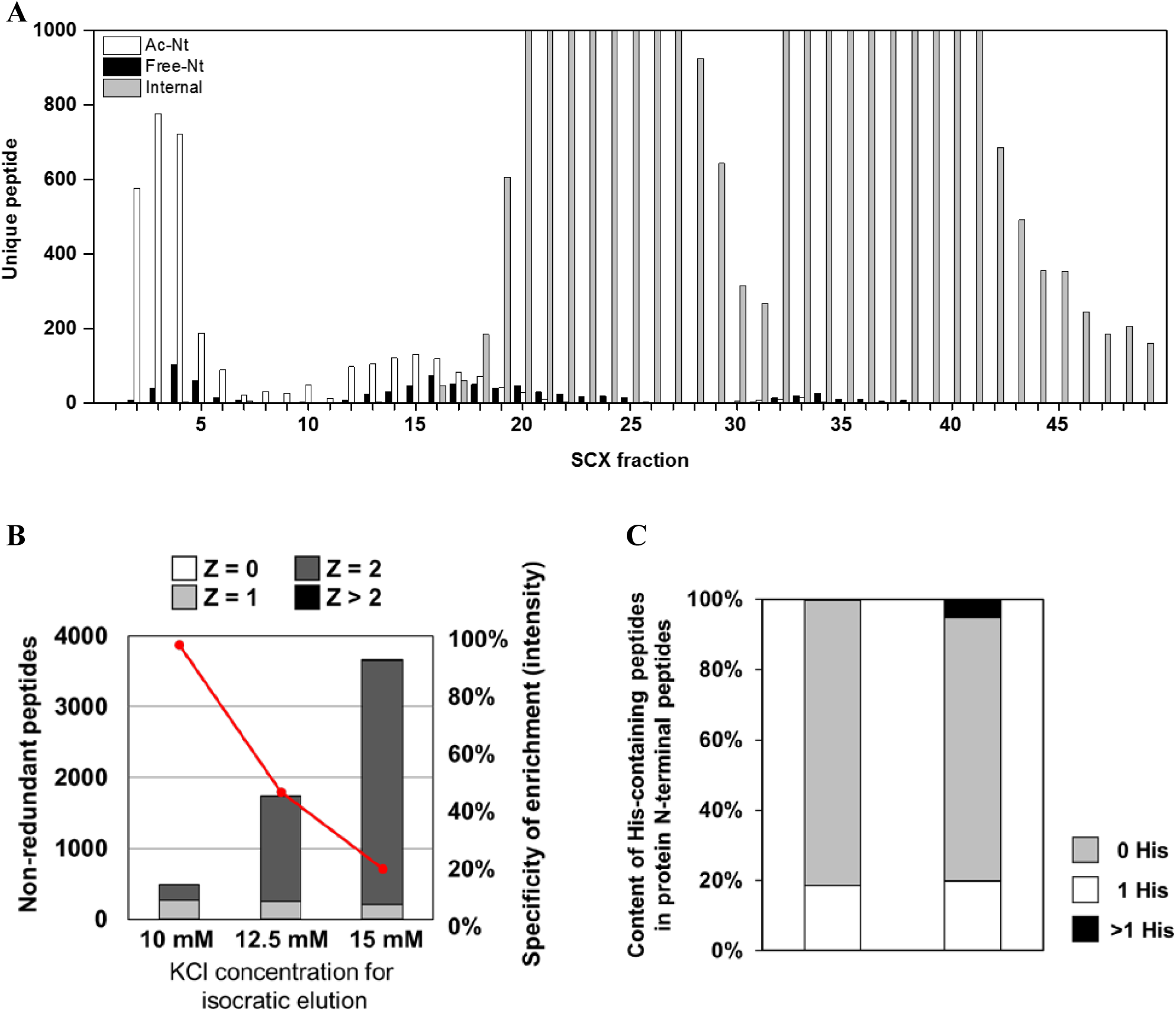
SCX elution profiles of TrypN-digested peptides. (A) SCX HPLC fractionation of different types of peptides in TrypN-digested HEK293T cell lysates using KCl salt gradient elution. In fractions 2-6, protein N-terminal peptides with Z = 0 and 1 (Z is the total of unmodified N-terminus and the number of basic amino acid residues) were observed, such as acetylated protein N-terminal peptides with or without a basic amino acid residue and unmodified protein N-terminal peptides without a basic amino acid residue. (B) SCX HPLC separation of TrypN-digested *E. coli* peptides under isocratic conditions with different KCl concentrations. The specificity in enriching protein N-terminal peptides and the numbers of peptides with different Z values are shown. (C) The content of His-containing peptides in protein N-terminal peptides. The experimental results were those obtained with 10 mM KCl isocratic elution, and the *in-silico* results were calculated from the *E. coli* K-12 MG1665 protein sequence database. Details of *in-silico* digestion are described in the supplemental methods.

### Optimization of CHAMP-SCX separation using TrypN-digested *E. coli* peptides

To optimize the elution conditions for CHAMP separation, we employed *E. coli* TrypN-digested peptides. Because bacterial proteins have less N-terminal modification than mammalian proteins, the bacterial sample was considered preferable to optimize the conditions for separating the protein N-terminal peptides with 2+ charge (peptides with an unmodified N-terminus and one His residue) from the internal peptides. Three SCX buffers with different KCl concentrations were used for isocratic elution for 30 min, and the enrichment efficiencies for protein N-terminal peptides were compared. An enrichment specificity of more than 97% was obtained with 10 mM KCl (Table 1). When buffers with higher KCl concentrations were used, more internal peptides were identified (Figure 2B). In the case of 10 mM KCI buffer, we identified 53 His-containing protein N-terminal peptides out of 270 non-redundant protein N-terminal peptides without missed cleavage from 20 μg of *E. coli* lysate (19.6%, Figure 2C). Among *in-silico* TrypN-digested peptides from the *E. coli* proteome, 20% of the protein N-terminal peptides contain one His, suggesting that our enrichment conditions have no bias in identifying His-containing protein N-terminal peptides. In other words, CHAMP-SCX chromatography was able to separate the protein N-terminal peptides from the internal peptides even in the most difficult cases where the unmodified protein N-terminal peptides contain an additional basic amino acid, such as His, Lys or Arg.

**Table 1.**
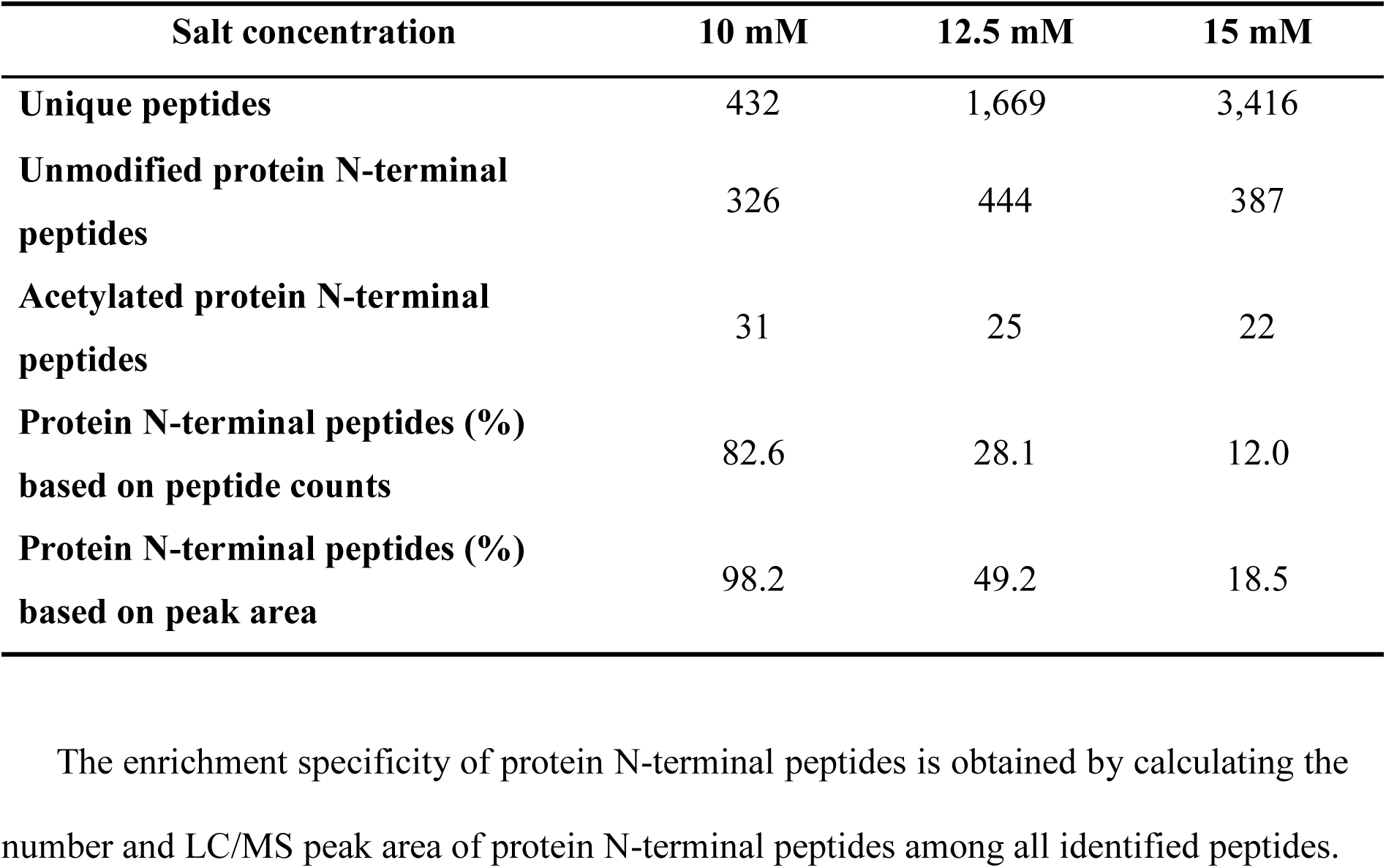
Enrichment of *E. coli* protein N-terminal peptides by SCX HPLC with isocratic elution at three different KCl concentrations.

| **Salt concentration** | **10 mM** | **12.5 mM** | **15 mM** |
| --- | --- | --- | --- |
| **Unique peptides** | 432 | 1,669 | 3,416 |
| **Unmodified protein N-terminal peptides** | 326 | 444 | 387 |
| **Acetylated protein N-terminal peptides** | 31 | 25 | 22 |
| **Protein N-terminal peptides (%) based on peptide counts** | 82.6 | 28.1 | 12.0 |
| **Protein N-terminal peptides (%) based on peak area** | 98.2 | 49.2 | 18.5 |

The enrichment specificity of protein N-terminal peptides is obtained by calculating the number and LC/MS peak area of protein N-terminal peptides among all identified peptides.

### HEK293T protein N-terminal peptide enrichment by CHAMP-SCX

The N-terminal peptides of His-containing proteins could be successfully separated from the internal peptides of human and bacterial samples by means of CHAMP-SCX under optimized elution conditions. To validate the applicability of this method to large-scale N-terminal proteomics, we performed triplicate analyses using HEK293T cells, which have been widely used in N-terminal proteomics (17, 18). Triplicate SCX fractionations using 10 mM KCl isocratic elution were done for TrypN-digested HEK293T peptides (80 μg each time), and we subjected one-fourth of the isolated peptides to nanoLC/MS/MS in triplicate (9 runs in total). Default parameters, such as the Swiss-Prot human protein sequence database, specific TrypN cleavage, and minimum peptide length of 7 amino acids, were applied for peptide identification by database search (Figure 3A). The results are shown in Figure 3B and Table 2. High correlations of peak areas of identified peptides were observed for intra- and interday preparation samples, (R^2^ = 0.95 and R^2^ = 0.75, 0.80, respectively). On average, we identified 1,550 unique acetylated and 200 unmodified protein N-terminal peptides from 20 μg of TrypN-digested HEK293T peptides in a single LC/MS/MS analysis. Contamination by internal peptides amounted to only 3% and 9% in peptide peak area and peptide number, respectively (Figure 3C, Table 2). Protein N-terminal peptides with missed cleavage were also enriched in the same elution, and 850 (∼50%) miscleaved unique N-terminal peptides were identified on average, improving the coverage of the N-terminome. We identified 1,640 acetylated, 106 partially acetylated and 167 unmodified non-redundant protein N-termini. Note that 1,600 additional neo-N-terminal peptides were identified when semi-specific cleavage at the N-terminus was allowed in the data processing, although our purpose in this study was not to find novel proteoforms, but to establish a novel approach for N-terminomics. Furthermore, to compare our results with two published N-terminome datasets for HEK293T human cells (17, 18), we re-analyzed those datasets under the same conditions without the use of their original customized database or non-specific cleavage. In terms of the contents of acetylated and unmodified protein N-terminal peptides, all three datasets provided identical results, whereas the content of internal peptides as well as the number of unique peptides varied depending on the approach and the sample amount (Supplementary Figure S4).

**Table 2.** Identification of protein N-terminal peptides from TrypN-digested HEK293T cells.

|  | **Replicate 1** | **Replicate 2** | **Replicate 3** | **Total** |
| --- | --- | --- | --- | --- |
| **Unmodified protein N-terminal peptides** | 199 ($\pm 3$) | 187($\pm 5$) | 197 ($\pm 3$) | 352 |
| **Acetylated protein N-terminal peptides** | 1,854 ($\pm 13$) | 1,301 ($\pm 18$) | 1,509 ($\pm 15$) | 2,666 |
| **Internal peptides** | 160 ($\pm 8$) | 147 ($\pm 3$) | 232 ($\pm 6$) | 433 |
| **N-term ratio (% , peptide counts)** | 92.8 ($\pm 0.3$) | 91.0 ($\pm 0.2$) | 88.0 ($\pm 0.3$) | |
| **N-term ratio (% , peptide area)** | 97.4 ($\pm 0.3$) | 98.0 ($\pm 0.5$) | 97.4 ($\pm 0.5$) | |
| **Unmodified protein groups** | 115($\pm 3$) | 100 ($\pm 4$) | 116 ($\pm 3$) | 167 |
| **Partially acetylated protein groups** | 36 ($\pm 2$) | 60 ($\pm 3$) | 50 ($\pm 2$) | 106 |
| **Acetylated protein groups** | 1,223 ($\pm 7$) | 1,000 ($\pm 9$) | 1,187 ($\pm 16$) | 1,640 |

Samples were prepared in triplicate (Replicates 1-3) and nanoLC/MS/MS of each sample was conducted in triplicate. Each number in the table is the average of triplicate measurements, and the total number is calculated after merging all results (n=9) and removing redundancy. The enrichment specificity of protein N-terminal peptides is obtained by calculating the number and peak area of protein N-terminal peptides among all identified peptides.

**Figure 3.**
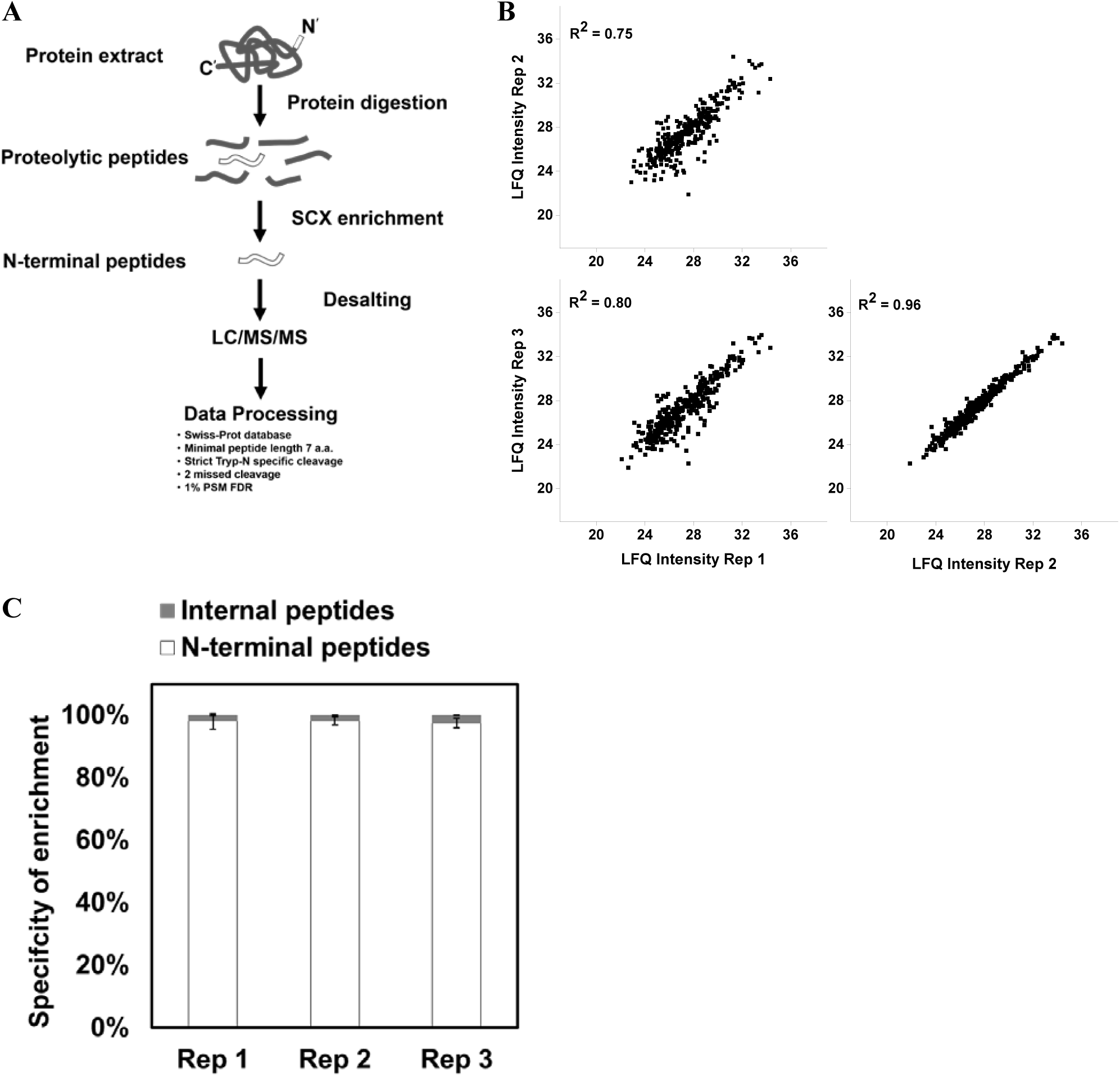
N-Terminal proteomics using CHAMP-SCX. (A) Workflow of TrypN/CHAMP-based N-terminal proteomics. For details, see Materials and Methods. Three technical replicates were conducted on Day 1 (Rep 1) and Day 2 (Rep 2 and Rep 3) to evaluate inter- and intraday reproducibility. (B) Reproducibility in quantifying peptide peak areas between three technical replicates. (C) Enrichment specificity for protein N-terminal peptides in three repeats.

In conclusion, we have succeeded in developing a new N-terminomics method that does not require chemical reactions. This simple and rapid approach is suitable for high-throughput screening with minimal sample amounts. Our TrypN/CHAMP-based N-terminomics can enrich protein N-terminal peptides without bias, including peptides containing basic amino acids, with or without N-terminal modifications. We believe CHAMP-SCX has great potential for expanding N-terminomics. Potential developments include deeper profiling with additional fractionation, the use of customized databases containing predicted protein N-termini, the replacement of HPLC with StageTips for CHAMP separation, and quantification by isotopic labeling.

## Abbreviations

CHAMP: charge-mediated position-selective enrichment
SCX: strong cation exchange chromatography
MS: mass spectrometry
COFRADIC: combined fractional diagonal chromatography
TAILS: terminal amine isotopic labeling of substrates
ChaFRADIC: charge-based fractional diagonal chromatography
HYTANE: hydrophobic tagging-assisted N-termini enrichment
Tris-HCl: 2-amino-2-(hydroxymethyl)-1,3-propanediol hydrochloride
SDC: sodium deoxycholate
SLS: sodium N-lauroylsarcosinate
TCEP: tris(2-carboxyethyl)phosphine
CAA: 2-chloroacetamide
ACN: acetonitrile,
TFA: trifluoroacetic acid
PTS: phase-transfer surfactants
StageTip: stop and go extraction tip

## ACKNOWLEDGMENTS

We would like to thank members of Department of Molecular & Cellular BioAnalysis for fruitful discussions. This work was supported by JST Strategic Basic Research Program CREST (No. 18070870) and JSPS Grant-in-Aid for Scientific Research No. 17H05667.

## DATA AVAILABILITY

All LC/MS/MS data that support the findings of this study have been deposited with the ProteomeXchange Consortium via the jPOST partner repository with the dataset identifier <u>JPST000422 (</u>PXD010551) (34).

